# Multivalent interaction induces phase separation and formation of more toxic aggregates of α-syn in a yeast model of Parkinson’s disease

**DOI:** 10.1101/2025.04.14.648683

**Authors:** Rajeev Jain, SK Sharavanakkumar, Krishnananda Chattopadhyay

## Abstract

The process of protein phase separation, particularly in the context of intrinsically disordered proteins, has been extensively studied for its implications in several neuro-degenerative diseases. Although the mechanism of protein phase separation and the involved molecular grammar have been well explored under in vitro conditions, the focus is now shifting towards developing more complex models of phase separation in order to mimic the biological systems more closely. Here, we studied the phase separation of alpha synuclein (α-syn), an intrinsically disordered protein whose aggregation is implicated in the pathology of Parkinson’s Disease, (PD) inside yeast cells (*Saccharomyces cerevisiae*). Using a positively charged polymer; polyethylenimine (PEI), which binds presumably at the negatively charged C-terminal domain of α-syn, we find that the aggregation of α-syn inside yeast can be modulated by at least two pathways: one involving phase separation and the second one without phase separation. We find further that these two pathways lead to varying fibril characteristics and toxicities. We believe that this model can be used as a quick and convenient system to screen novel and repurposed small molecules against toxic protein droplets.

## Introduction

Parkinson’s disease (PD) is the second most common neuro-degenerative disease after Alzheimer’s Disease (AD). PD is characterised by motor disorders, which are typically manifested by tremors, muscular rigidity, bradykinesia among other symptoms (1). It is believed that the pathology of PD is caused by the loss of dopaminergic neurons in substantia nigra pars compacta (SNPC), and the appearance of cytoplasmic inclusions, known as Lewy bodies and Lewy neuritis. Accumulation of cytoplasmic inclusions causes degeneration of dopaminergic neurons, resulting in the deficiency in dopamine neurotransmitter as the disease progresses (1). Alpha Synuclein (α-syn), a 140 amino acid long protein is considered to be one of the important players in PD, which is known to be found in Lewy bodies in the filamentous form (2). This intrinsically disordered protein (IDP) consists of three domains, a positively charged N-terminal domain (NTD) spanning residue 1-60, a central hydrophobic non-amyloid-β component (NAC) spanning residue 61-95 and a negatively charged C-terminal domain (CTD) spanning residue 96-140. Inside neurons, it primarily localizes in the presynaptic part both as soluble and membrane bound forms. Although the exact physiological function of α-syn is not yet determined, various studies suggest that α-syn is responsible for vesicles trafficking during neurotransmitter release (3).

The fibrillation of α-syn follows a nucleation dependent pathway of aggregation, through which the protein first undergoes a primary nucleation phase and few native monomeric protein may misfold and combines to form a nucleus. This step may then be converted into oligomeric seeds by combining with more monomeric units, which can then be followed by the formation of fibrillar aggregates (4, 5). Recent studies suggest that in addition to the classical pathway which was described above, the protein under stressed conditions can also undergo liquid-liquid phase separation (LLPS) to form protein droplets. The protein droplets, which are liquid like but without any membrane, can subsequently undergo liquid to solid transition forming fibrillar aggregates. These liquid droplets can also be formed inside cells which subsequently turns to solid aggresomes (6). It has been demonstrated further that apart from the neurodegeneration, protein phase separation is involved in various other cellular functions, like transcription, stress granule assembly, signal transduction, immune signalling, heterochromatin formation, and also in infectious disease progressions (7). SARS-CoV-2 N protein forms phase separated bodies with RNA to form ribonucleoprotein which promotes genome packaging and virion assembly (8). In addition, leishmania protein KMP-11 undergoes phase separation thus promoting infection by facilitating parasitic entry into host cell (9).

Hydrophobicity of proteins and weak multivalent interactions between protein monomers facilitates a number of homotypic and heterotypic interactions with the same protein molecules or other molecules inside the cell promoting phase separation (10). Other modulators which contribute to phase separation behaviour includes local environmental factors or stress conditions, like the change in pH, temperature, salt concentration, (11) etc and/or the presence of specific chemicals (12, 13), metal ions (14), small molecules (15) and metabolites (16, 17). *In-vitro* phase separation of α-syn can be modulated by the presence of molecular crowders like polyethylene glycol (PEG) and by altering pH and salt concentrations (11).

Cellular models help in better understanding of basic mechanism and pathways underlying neurodegenerative diseases, such as proteotoxicity, disease progression, mitochondrial dysfunction, transcriptional dysregulation, trafficking defects, proteasomal dysfunction, etc (18). *Saccharomyces cerevisiae* or budding yeast is one such well validated model which is extensively used in different mechanistic studies involving the biology and human diseases (19). One reason could be that several mechanisms and pathways are extremely well conserved between human and yeast (18). Although the classical aggregation pathway (which does not mediate through the phase separation) has been extensively studied in yeast, the phase separation of α-syn protein is not shown in yeast until now. Here in this study, we for the first time, have established a phase separation model of α-syn protein inside yeast using a positively charged polymer, polyethylenimine (PEI). With the help of various biophysical, microscopy and biochemical assays we show that PEI not only helps α-syn to form liquid condensates presumably by modulating the long-range electrostatic interactions between the monomers, it also drives the short-range hydrophobic interactions stabilizing the droplets formed. We also found that the addition of PEI enhances the rate of fibrillation and produces aggregates having distinct morphology which are more toxic than the classical aggregation pathway. The droplets inside yeast are further characterized by monitoring fluorescence lifetime measurements using TD-FILM. We believe that this yeast LLPS model of PD will help in better understanding of disease progression involving various machineries and pathways thus allowing researchers to develop condensate targeting novel molecules.

## Materials and Methods

For the purification of recombinant α-syn protein, tris base, sodium chloride, ammonium sulphate, streptomycin sulphate, ethylenediaminetetraacetic acid, phenylmethanesulfonylfluoride (PMSF), isopropyl β-D-galactopyranoside (IPTG), sodium phosphate dibasic (Na_2_HPO_4_) and sodium phosphate monobasic (NaH_2_PO_4_) were used and purchased from Sigma Aldrich, St. Louis, USA. Thioflavin T, sodium dodecyl sulphate (SDS), glycine, ammonium persulfate, N, N, N′, N′ -tetramethylethylenediamine were purchased from Sigma Aldrich. Polyethylenimine (PEI) was purchased from Sigma Aldrich, St. Louis, USA whereas 1,6-Hexanediol was purchased from Himedia, Thane, India. All chemicals used in this study were of highest grade.

### Molecular docking analysis of the binding between α-syn and PEI

#### Preparation of receptor and ligand

The 3D structure of α-syn was retrieved from the Protein Data Bank (PDB ID: 1XQ8). The structure was energy minimized using GROMOS96 force field available with the Swiss-PDB Viewer (20) and used as the receptor molecule, while the 3D structure of branched PEI was drawn using MarvinSketch 20.9.0 (http://www.chemaxon.com) and exported in MD mol (.sdf) format. Only four monomeric units were used in the polymeric ligand, PEI considering the constraint on acceptable number of rotatable bonds in the AutoDock algorithm, the ligand was further energy minimized using AMBER ff14SB force field present in Chimera 1.17.1 (21) and then converted to PDB format, acceptable in AutoDock using Open Babel 2.4.1 (22).

#### Docking simulations

AutoDock Tools (ADT) (23) were used to prepare the receptor and ligand molecules. Polar hydrogens, Kollman charges and AD4 type of atoms were added to the receptor molecule, while ligands were added with Gasteiger charges and maximum numbers of active torsions were maintained. AutoGrid was used to prepare a grid map. Because of the lack of information on specific binding sites in α-syn monomer with PEI or any polymeric ligands, grid box was set covering the whole protein structure and docking was performed to avoid any bias in obtaining the structural domain which interacts with PEI. For α-syn a grid box of 62 X 120 X 50, centered on X, Y, Z of 227.401, -3.874, -11.978 covering the whole receptor structure. A grid spacing of 0.96 Å was maintained for the docking procedures. Lamarckian Genetic Algorithm (LGA) was used for performing molecular docking, keeping the receptor molecule rigid throughout the docking simulation. The population size was set to 150 and the individuals were initialized randomly. Maximum number of energy evaluations was set to 2500000. Rest of the docking parameters was set to default values. Ten different poses were generated for each ligand and scored using AutoDock 4.2 scoring functions and were ranked according to their docked energy. Dockings that resulted in the lowest docking energy with higher number of H-bonds were selected. For post docking analysis PyMOL (https://www.pymol.org) software was used.

#### Expression and purification of α-syn

Wild type recombinant α-syn protein was expressed in *E. coli* BL21 (DE3) strain harbouring pRK172 plasmid. When the culture reached an OD_600_ of 0.6-0.8, the protein was induced by adding 1 mM IPTG and incubating for 4 h at 37 °C under stirring at 180 rpm. Cells were harvested by centrifugation. The protein was purified using non-chromatographic method, which has been reported previously (24). Briefly, cell pellet was re-suspended in Lysis buffer containing 50 mM Tris, 10 mM EDTA and 150 mM NaCl (pH 8.0) followed by boiling using a water bath for 10 min. 1 mM PMSF was added prior to boiling. Cell debris and supernatant were separated by centrifugation under 15000*g* for 20 min at 4 °C. After collecting the supernatant in a fresh tube streptomycin sulphate (136 µL of 10% solution/ml of supernatant) and glacial acetic acid (228 µL/ml of supernatant) were added. The pellet and supernatant were again separated by centrifugation at 15000*g* for 20 min at 4 °C. The supernatant was taken in a beaker and precipitated by adding small volumes of ammonium sulphate to maintain 50% saturation (calculated from EnCor Biotechnology Inc.) with constant stirring at 4 °C. Precipitated protein was separated by centrifugation at 15000*g* for 20 min at 4 °C and protein pellet obtained was washed once with 50% ammonium sulphate solution followed by centrifugation under 15000*g* for 20 min at 4 °C. The washed pellet was re-suspended in equal volume of 100 mM ammonium acetate and ethanol followed by centrifugation at 15000*g* for 20 min at 4 °C. The above step was repeated twice. Finally, the protein pellet obtained was dissolved in 20 mM sodium phosphate buffer (NaP), pH 7.4 and dialysed overnight using a SnakeSkin dialysis tubing (10 kDa MWCO) against NaP at 4 °C. The dialysed protein was collected as small fractions in microcentrifuge tubes, flash froze under liquid nitrogen, lyophilized and stored at -80. For sample preparation of all *in-vitro* experiments, lyophilized α-syn protein was dissolved in 20 mM NaP buffer (pH 7.4) followed by centrifugation under 15000*g* for 1 h at 4 °C for removal of any trace amount oligomeric forms that may form during purification and storage (25). Pure α-syn obtained this way was analysed by SDS-PAGE (15%) (26) and the bands appeared at approx. 15 kDa (Supplementary Fig. 4a) which corresponds well with the theoretical molecular weight of wild type recombinant α-syn proteins respectively. Mass spectrometric analysis also confirmed the presence of 14.5 kDa monomeric α-syn protein (Supplementary Fig. 4b).

#### Fluorescence labelling of α-syn protein with Alexa Fluor^TM^ 488 C_5_ maleimide

Since wild type recombinant α-syn protein does not contain any cysteine amino acid in the peptide chain, we introduced a cysteine mutation (G132C) at the C-terminal of the protein. By mutating glycine by cysteine at 132nd position, the physicochemical as well as aggregation propensity was found to be identical to that of wild type protein (27, 28). This G132C mutant was then labelled with Alexa Fluor^TM^ 488 C_5_ Maleimide dye (ThermoFisher Scientific, USA) following the manufacturer’s protocol.

#### PEI-α-syn binding analysis

In order to study the binding of PEI with α-syn, steady state fluorescence spectroscopy was utilized. Briefly, titration was performed with addition of an increasing concentration of PEI (0.0267 nM – 2.67 µM) onto a fixed amount of α-syn protein (20 µM) doped with 1% of labelled α-syn protein. Upon binding to increasing PEI concentration there was a change in fluorescence intensity at emission wavelength of 516 nm (excitation wavelength of 493 nm). The measurements were recorded using a quartz cuvette of 1 cm path length using a PTI spectro fluorimeter (Kyoto, Japan). The fluorescence values obtained were plotted against the PEI concentration and was fit into a nonlinear one site-total binding equation approaching saturation for getting the dissociation constant (K_d_) value.

#### Turbidity assay

For measuring the change in turbidity of the solutions during non-LLPS or LLPS conditions, UV-visible spectra of α-syn protein (100 µM) incubated in absence or presence of PEI were recorded after incubating the samples for 30 min at 350 nm using a UV-visible spectrophotometer (Agilent Cary 60 UV-Vis, Santa Clara, California, U.S). Final picture of the cuvettes was also taken against a white background with a black strip for the visual detection of turbidity of the solution.

#### Confocal microscopy

All imaging studies were carried out using a Zeiss LSM 980 inverted microscope equipped with a Plan-Apochromat 63X/1.4 NA oil immersion objective. 20 µM α-syn protein doped with 1% of labelled protein without and with PEI were incubated for 15 min. Samples were cast on a clean glass slide and images were taken using a 488 nm laser. Images were collected and processed using the Zen Blue software provided by Carl Zeiss Inc, Oberkochen, Germany. Circularity of the droplets were calculated using imageJ software.

#### Thioflavin-T aggregation assay

Aggregation of α-syn during non-LLPS or LLPS conditions was monitored using the emission of the Thioflavin T dye (29). Briefly, 200 µM of freshly dissolved lyophilized α-syn in NaP buffer (pH 7.4) was incubated without and with PEI under constant shaking condition at 200 rpm and 37 °C. At different time interval, suitable aliquots were withdrawn from each sample, diluted to 400 µL volume in NaP buffer and the dye was added maintaining the protein to ThT ratio of 1:3. Steady state ThT fluorescence was measured at an excitation wavelength of 450 nm and emission spectra between 460 to 500 nm were recorded using a PTI fluorometer (Kyoto, Japan). Blank solutions containing only the buffer and ThT were also prepared. Fluorescence spectra for each blank samples were also recorded and subtracted from those of the samples. The normalized fluorescence values were plotted against time of incubation and was fit into a Boltzmann sigmoidal equation to obtain an aggregation specific curve.

#### Atomic force microscopy

At different time intervals, 10 µL aliquots from the same samples used in ThT aggregation assay were withdrawn. Aliquots were adsorbed onto a freshly peeled mica sheet followed by drying in a vacuum desiccator for 2 h. Gentle washing of dried samples were performed with 200 µL milliQ water followed by overnight drying in a vacuum desiccator. The adsorbed α-syn protein was imaged using Acoustic Alternative Current (AAC) or tapping mode by Agilent Technologies Picoplus AFM 5500 having a 9 µm piezoscanner with a scan frequency of 1.5 Hz. Microfabricated silicon cantilevers of 225 μm in length with a nominal spring force constant of 21–98 N/m (obtained from Nano Sensors). The cantilever oscillation frequency was tuned to the resonance frequency. The cantilever resonance frequency was 150–300 kHz. The images (256 pixels × 256 pixels) were captured with a scan size of between 0.5 and 5 μm at a scan speed of 0.5 lines/s. Images were processed and analysed by flattening using PicoView software (Agilent Technologies, USA).

#### Gel Partitioning and apparent partition Gibb’s free energy

For measuring gel partitioning 100 µM of freshly dissolved lyophilized α-syn in NaP buffer (pH 7.4) was incubated with different concentrations of PEI under constant shaking conditions at 200 rpm and 37 °C. After 24 h, the solution was gently centrifuged at 4000 rpm for 10 min to separate dense turbid phase (D) and light solution phase (L). Aliquots from both phases were loaded onto 15% SDS-PAGE (26). Gel partitioning was calculated by measuring the ratio of band intensity of dense phase to that of light phase. From this gel partitioning ratio, apparent partition Gibb’s free energy was calculated using the following equation:

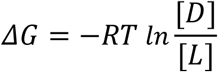

where R is ideal gas constant, T is temperature in kelvin, [D] and [L] are the band intensities of dense phase and light phase respectively. More the negative value of ΔG more spontaneously phase separation will occur.

#### Circular dichroism (CD)

Change in the secondary structure was monitored by far UV-CD using a MOS-500 spectropolarimeter (BioLogic, Seyssinet-Pariset, France) keeping the scan speed at 50 nm/min, integration time at 1 s, and number of acquisitions equal to 3. A CD cuvette with 1 mm path length was used for the far UV CD measurements. These CD measurements were carried out between 195 and 250 nm using the permissible HT voltage to obtain an optimum signal-to-noise ratio. At different time intervals, 10 µL of the incubated samples of α-syn protein with different PEI concentrations were aliquot to a final concentration of 10 µM before the final measurements.

#### Yeast strain, growth and expression of α-syn

*Saccharomyces cerevisiae* W303-1B cells (*MATα leu2-3,112 trp1-1 can1-100 ura3-1 ade2-1 his3-11,15*) were transformed with a p426-GAL1-*WTα-syn-GFP* vector (30). Transformed cells were grown in SC-URA + 2% dextrose (SD medium) at 30 °C, 200 rpm till OD_600_ 0.8 –1.0 and protein expression was induced with SC-URA + 2% galactose (SGal medium) in the absence or presence of PEI. At different time interval, expression of α-syn was monitored by fluorescence microscopy (BX41, Olympus, Tokyo, Japan). For the dissolution of droplets aliquots at different time intervals were incubated with 10% of 1,6-Hexanediol for 1 h. After 1 h cells were washed twice with PBS and imaged by fluorescence microscope.

For mitochondrial morphology analysis, at different time intervals aliquots of induced cells without or with PEI were further processed with MitoTracker^TM^ Red FM dye (ThermoFisher Scientific, USA) following the manufacturer’s protocol and mitochondrial morphology was visualized under fluorescence microscopy (BX41, Olympus).

#### Fluorescence recovery after photo-bleaching (FRAP)

Induced yeast cells expressing α-syn protein without or with PEI were aliquoted and subjected to fluorescence recovery after photobleaching (FRAP) analysis selecting a region of interest (ROI) of constant radius inside the foci. ROI was bleached using 488 nm laser at 100% intensity using Zeiss LSM 980 inverted microscope equipped with a Plan-Apochromat 63X/1.4 NA oil immersion objective and subsequent recovery of the bleached area was recorded with 488 nm laser till 90 s. Recovered fluorescent data was analysed using Zen Blue software (Zeiss). % fluorescence recovery for different samples were calculated from the initial pre-bleached, bleached and final recovered intensities and plotted to conclude the nature of the foci.

#### TD-FLIM analysis of α-syn protein inside yeast

Induced yeast cells in absence and presence of PEI were aliquot at different time intervals and washed twice with PBS. Cells were cast on cleaned glass slide and imaged with 488 nm laser using an ISS Alba FFS/FLIM Confocal Microscope (Champaign, IL, USA) equipped with a 60X water-immersion objective (NA 1.2). The time-resolved fluorescence measurements inside yeast cells were made using a time-correlated single photon counting (TCSPC) setup of ISS Alba FFS/FLIM Confocal Microscope (Champaign, IL, USA). The samples were excited using a 488 nm QuixX picosecond pulsed laser made by Omicron-Laserage Laserprodukte GmbH (Germany). The repetition rate of the laser was set to be 20 MHz. The fluorescence emission was detected by a single photon avalanche diode (SPAD) detector (SPD-100-CTC by Micro Photon Devices) after the 530/43-nm band-pass filter (Semrock, NY, USA). TD-FLIM data was analysed using the ISS 64-bit VistaVision software. It allows single or multiexponential curve fittings on a pixel-by-pixel basis using a weighted least-squares numerical approach (31–33). The double-exponential model was used for fitting the lifetime data of diffused, condensed and aggregated α-syn protein inside yeast with the instrument response function, estimated by taking the first derivative of the rising of the decay.

#### Cytotoxicity assays

To measure the toxicity of PEI on yeast cells, serial dilution spot assay was performed (34). Briefly yeast cells were grown in SD media at 30°C, 200 rpm till OD_600_ value of 1.0. Cells were harvested by centrifugation, washed twice with sterile milliQ water, treated with different concentrations of PEI and incubated for 48 h. After 48 h, cells were serially diluted in ten-fold increments in PBS. 5 µL from each dilution was spotted onto SD agar plates. Yeast growth was visualized after incubation for 24-36 h at 30 °C.

For measuring the colony forming units (CFU) (35), yeast cells were grown in SD media at 30 °C, 200 rpm till OD_600_ value of 0.8 – 1.0, induced with SGal medium and PEI was added during induction. At different time interval aliquots of each sample were taken, cells were harvested by centrifugation, washed twice with sterile milliQ water, diluted 500-fold with sterile milliQ water and spread onto SD agar plate. Colonies were allowed to form for 48 h and colony forming units (CFU) were recorded after incubating the plates at 30 °C.

Cells were visualized under fluorescence microscope and cells positive for red fluorescence were scored.

#### Lysis of yeast cells

Yeast cells were lysed using mechanical glass beads disruption method (36). Cells were grown in SD medium at 30 °C, 200 rpm till OD_600_ 0.8 – 1.0 and protein expression was induced with SGal medium in the absence or presence of PEI. After 8 h of incubation, cells were harvested by centrifugation, washed twice with sterile milliQ water and the cell pellet was kept in -20 °C overnight. Cells were then re-suspended in 500 μl lysis buffer containing Tris HCl (50 mM), NaCl (150 mM), DTT (2 mM), glycerol (10 %), Triton X-100 (1 %), (pH 7.5) supplemented with 1 mM PMSF. Cells were disrupted using acid washed glass beads by vortexing 10 times for 1 min with intermittent rests of 30 sec on ice. After lysis, the tubes containing the lysate were set aside for 1 h for the glass beads and cell debris to settle down. The lysate was then transferred to another centrifuge tube and was centrifuged at 15000 g for 1 h. Supernatant was collected in a fresh tube. Protein estimation was done in the resulting supernatant by dye binding method (37) using bovine serum albumin as the standard protein.

#### Western blot analysis

Lysed supernatant 200 µg/ml protein was run on 12% SDS-PAGE (26) using vertical electrophoresis system (BioRad) under constant current. After carrying out SDS-PAGE, the proteins on the gel were transferred to a nitrocellulose (NC) membrane (0.45 μm) with transfer buffer [25 mM Tris, 20 mM glycine, and 10 % (v/v) methanol, pH 8.3] at 20 V, 3 A for 30 min on a semi dry blotting assembly (BioRad). In order to block nonspecific binding, the nitrocellulose membrane was kept for 3 h in 5 % (w/v) BSA solution. After completion of blocking, the membrane was washed with TBST (Tris-buffered saline pH 7.2 and 0.1 % v/v Tween 20) and TBS thrice, for 5 min each and the membrane was incubated with rabbit anti-GFP (1:2500) overnight at 4 °C. After washing the membrane with TBST and TBS thrice, goat anti-rabbit HRP conjugated secondary antibody (1:2500) was added to it and incubated for 1 h at room temperature. Finally, the blot was washed with TBST and TBS and developed using SuperSignal West Pico PLUS chemiluminescent substrate (ThermoFisher Scientific, USA) with an iBright imaging System (ThermoFisher Scientific, USA). Band intensities were estimated using imageJ.

#### Flow cytometry analysis

CellRox^TM^ Deep Red (ThermoFisher Scientific, USA), a far-red fluorescent dye (absorption/emission maxima at ∼644/665 nm) which is non-fluorescent while in a reduced state and becomes fluorescent upon oxidation by reactive oxygen species (ROS) was used for reactive oxygen species analysis. Yeast cells were grown on SGal medium in presence and absence of PEI at 30 °C. At different time interval yeast cells were harvested by centrifugation, washed twice with sterile milliQ water and further processed with CellRox^TM^ Deep Red dye (ThermoFisher Scientific, USA) following the manufacturer’s protocol. At least 10,000 cells (events) were collected for each specimen and analysis was conducted on viable cells using a BD LSRFortessa™ cell analyser flow cytometer (BD Biosciences, New Jersey, U.S.).

Similarly for mitochondrial morphology analysis, yeast cells were grown on SGal medium in presence and absence of PEI at 30 °C. After 8 h yeast cells were harvested by centrifugation, washed twice with sterile milliQ water and were further processed with MitoTracker^TM^ Red FM dye (ThermoFisher Scientific, USA) following the manufacturer’s protocol. Cell-permeant MitoTracker^TM^ Red FM probe stain active mitochondria in live cells (absorption/emission maxima at ∼581/644 nm). At least 50,000 cells (events) were collected for each specimen and analysis was conducted on viable cells using a BD LSRFortessa™ cell analyser flow cytometer (BD Biosciences, New Jersey, U.S.).

For the mitochondrial ROS measurement, MitoSOX^TM^ Red dye was used. MitoSOX^TM^ Red superoxide indicators are novel fluorogenic dyes specifically targeted to mitochondria in live cells. Oxidation of the MitoSOX^TM^ reagent by mitochondrial superoxide produces bright red fluorescence. Briefly yeast cells were grown on SGal medium in presence and absence of PEI at 30 °C. After 8 h yeast cells were harvested by centrifugation, washed twice with sterile milliQ water and were further processed with MitoSOX^TM^ Red dye (ThermoFisher Scientific, USA) following the manufacturer’s protocol (absorption/emission maxima at ∼396/610 nm using custom filter set). At least 50,000 cells (events) were collected for each specimen and analysis was conducted on viable cells using a BD LSRFortessa™ cell analyser flow cytometer (BD Biosciences, New Jersey, U.S.).

#### Dot blot analysis

Yeast cells induced under Non-LLPS and LLPS conditions were grown for 8 h. Cells were harvested, lysed and clarified supernatant obtained were subjected to dot blot analysis performed in a 96 well format using dot blot apparatus (Cleaver Scientific, UK) using vacuum to transfer samples to nitrocellulose membrane pre-wetted with TBS buffer. In order to block nonspecific binding, the nitrocellulose membrane was kept for 3 h in 5 % (w/v) BSA solution. After completion of blocking, the membrane was washed with TBST (Tris-buffered saline pH 7.2 and 0.1 % v/v Tween 20) and TBS thrice, for 5 min each. For monomeric and fibrillar species analysis, membranes were incubated with rabbit anti-α-syn antibody (ab138501, abcam, Cambridge, UK) (1:10,000) and rabbit anti-α-syn aggregate antibody (ab209538, abcam, Cambridge, UK) (1:2000) respectively for overnight at 4 °C. After washing the membrane with TBST and TBS thrice, goat anti-rabbit HRP conjugated secondary antibody (31460, ThermoFisher Scientific, USA) (1:10,000) was added to the membranes and incubated for 1 h at room temperature. Finally, the blots were washed with TBST and TBS and developed using SuperSignal West Pico PLUS chemiluminescent substrate (ThermoFisher Scientific, USA) with a iBright imaging System (ThermoFisher Scientific, USA). Band intensities were estimated using imageJ.

#### Limited proteolysis using Proteinase K (PK)

Proteolytic fingerprinting of different fibrils was done by subjecting them to limited proteolysis by Proteinase K (PK) followed by running the digested samples on SDS-PAGE (26). Characteristic band patterns with positions and relative intensity were obtained on digestion depending on the epitopes of the protein accessible to the enzyme. 200 µM α-syn protein was allowed to form fibrils under non-LLPS and LLPS conditions by incubating for 96 h under constant agitation. Fibrils formed after incubation was pelleted down, washed with NaP buffer and passed through 1000 kDa cut-off spin column (Amicon Ultra-0.5 Centrifugal Filter Unit). The residual volume retained in the spin column contains fibrils >100 kDa. Equal volume of α-syn fibrils formed under non-LLPS and LLPS conditions were mixed with 0, 0.01, 0.1 and 1 μg/ml of PK (20 mg/mL) in NaP to a final volume of 20 μl, and incubated at 37°C for 30 min. PK digestion was stopped with 1 mM PMSF. Reaction samples were boiled with SDS loading buffer for 10 min at 95 °C. Samples were resolved onto 15% SDS gel to obtain the proteolytic fingerprints.

#### Data plotting and statistical analysis

All experimental data were plotted and fit using GraphPad Prism 8 software. Statistical data analyses were performed using t-test (wherever required), where ns: nonsignificant, *: p ≤ 0.05, **: p ≤ 0.01, ***: p ≤ 0.001, and ****: p ≤ 0.0001.

## Results

### *In-silico* analyses of α-syn

Supplementary Fig. 1a shows the sequence map of the protein α-syn (P37840), indicating the presence of a negatively charged C-terminal, an amphipathic N-terminal and a hydrophobic NAC domain. Intrinsic disorder propensity analysis of the α-syn using IUPred2A (https://iupred2a.elte.hu/) algorithm (38) suggests that the residues (96–140) covering the C-terminus contain low-complexity domains (LCD) (Supplementary Fig. 1b). Further analysis using FuzDrop (https://fuzdrop.bio.unipd.it) algorithm (39) predicts that this protein would undergo LLPS (Supplementary Fig. 1c). A comparison between these two tools clearly predicts that the propensity of this protein towards LLPS formation comes from the C-terminal through the involvement of the LCD regions.

### α-syn phase separates inside yeast

To investigate if α-syn undergoes LLPS inside *S. cerevisiae* cells, we grew yeast cells harbouring plasmid for α-syn as outlined in detail in the methods section. It is known that in yeast, α-syn expression and aggregation occurs by localization to membrane during initial hours of induction followed by the formation of intracellular foci/puncta in cytoplasm during 6-8 h of induction (40). We observed similar pattern of membrane localization of α-syn and formation of intracellular foci after 6 h of protein induction (Fig. 1a, top panel). As shown in previous literature, we also observed the formation of matured aggregates after 24 h incubation (Fig. 1a, top panel). In the presence of 5mM guanidinium hydrochloride, which was shown earlier to inhibit Hsp104 activity inside yeast (41), we found no change in the α-syn aggregation pattern (Supplementary Fig. 2a). The addition of 1-6 hexanediol, which has been shown to dissolve liquid like droplets formed due to weak hydrophobic interactions (42), did not show any effect (Fig. 1b, top panel). When we studied these foci/puncta using Fluorescence Recovery After Photobleaching (FRAP), we found about 46% recovery at 4 h, and the percentage of recovery did not change significantly after 8 h incubation (Fig. 1c).

Subsequently, we studied the effect of an anionic (heparin) and a cationic (PEI) agent to investigate if these influence the aggregation pattern of α-syn inside yeast. We chose heparin as it was shown earlier to facilitate aggregation and induce the formation of liquid like droplets in different protein systems (43–45). However, we did not see any effect of heparin when we carried out a systematic time dependence analysis (Supplementary Fig. 2b). PEI, on the other hand, has been shown earlier to alter conformation and aggregation kinetics of α-syn (46). In this case, when we added 0.026 µM or above concentration of PEI (majority of the experiments were done here in the presence of 0.133 µM), small puncta of α-syn appeared near the plasma membrane after about 2 h of incubation, which became more prominent after 4 h (Fig. 1a, bottom panel). Unlike in the case of untreated cells where 1,6 hexanediol did not have any effect, the addition of 10% 1,6 hexanediol dissolved these puncta, which formed on PEI treatment after 2 or 4 hrs incubation (Fig. 1b, bottom panel). Using previous literature (42), we suggest that while the puncta formed in the absence of PEI is more solid like (no effect of 1,6 hexanediol), those formed in the presence of PEI has liquid like behaviour (strong 1,6 hexanediol sensitivity). We confirm this by carrying out FRAP experiments, which showed 76% recovery for the puncta formed in the presence of PEI after 4 hours incubation (high recovery suggest liquid like behaviour as suggested before). It may be recalled that, in the absence of PEI, the percentage was less than 50% (suggesting less liquid like and more solid like behaviour). The percentage of fluorescence recovery decreased significantly when incubated in the presence of PEI beyond 8 hours. As a matter of fact, for both in the absence and presence of PEI, after 8 hours the recovery was less than 10% (Fig. 1c). We believe that, for both in the absence and presence of PEI), longer time incubation (8 hours and beyond) results in solid like aggregates. A comparison between the recoveries between the 8 hours data in the absence and presence of PEI seems to suggest a higher recovery in the absence of PEI (Figure 1c).

**Figure 1:**
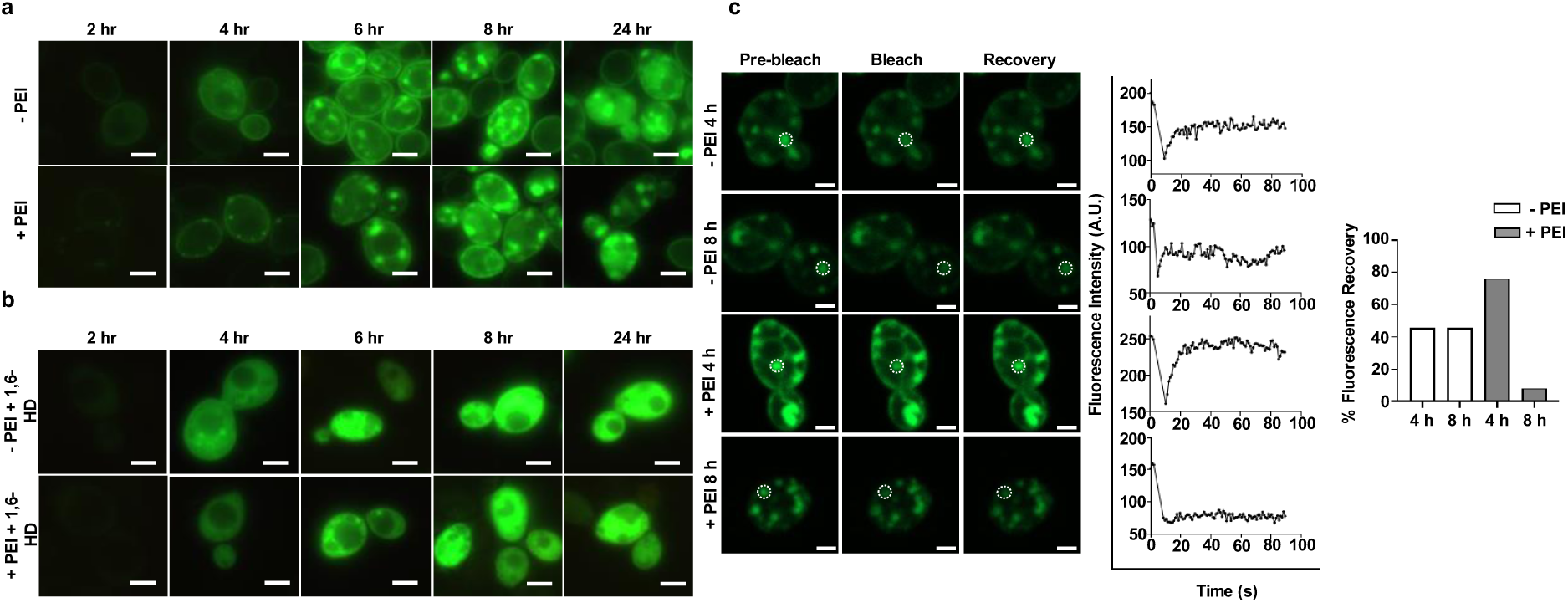
α-syn phase separates inside yeast: **a)** Fluorescence imaging of S. cerevisiae cells expressing WT α-syn-GFP in the absence and presence of PEI for different time intervals **b)** Fluorescence imaging of S. cerevisiae cells (treated without or with PEI) after treatment with 1,6-HD (10%) for 1 h [Bar = 5 μm; magnification = 100× (oil immersion objective)] **c)** Fluorescence recovery after photobleaching (FRAP) analysis of untreated and PEI treated cells at different time intervals showing % fluorescence recovery of the droplets inside yeast after photobleaching [Bar = 2 μm; magnification = 100× (oil immersion objective)]

The experiments discussed above thus indicated that in the presence of PEI, the aggregation of α-syn occurred through the formation of a liquid like state (phase separation pathway) before the formation of the final solid like aggregates. In contrast, in the absence of PEI, the formation of the final solid like state was direct without the formation of the liquid droplets like intermediates (non-phase separation pathway). To get further insights into these two pathways inside yeast, we carried out Fluorescence Lifetime Imaging Microscopy (FLIM) imaging. Fluorescence lifetime is an intrinsic property of the fluorophore, which has been shown to depend on local microenvironments, and not on the concentrations (33). Induced yeast cells under phase separation (in the presence of PEI) and non-phase separation (in the absence of PEI) pathways were imaged and analysed as described in methods section. The fluorescence lifetime of GFP tagged proteins typically vary between 2-5nsec (47), and for both pathways studied here, at the first time point (time 2 hour) the lifetime distributions provided a maximum at around 2.4 nsec which matches well with the previously reported values of GFP proteins (48). We found that the earlier time points (time points 4 hrs and 6 hrs), the lifetime distributions were wider for the non-phase separation pathway (Fig. 2). While it is possible that the process of phase separation somehow makes the aggregation pathways more homogeneous by entrapping different small molecules oligomers inside the droplets, this hypothesis requires further characterization and experimental validations. At the later time points, the lifetime distributions were similar in characteristics for both conditions, and similar mean values were obtained in both cases (Fig. 2). It may be noted that the mean lifetime values at the later time points were significantly less (1.5 nsec to 1.8 nsec) for both conditions, which is a reported characteristic of late-stage solid like aggregates of different proteins. Reports also suggests that difference in lifetime between late-stage aggregates of two different pathway under study have different characteristic or population of fibrils (49).

**Figure 2:**
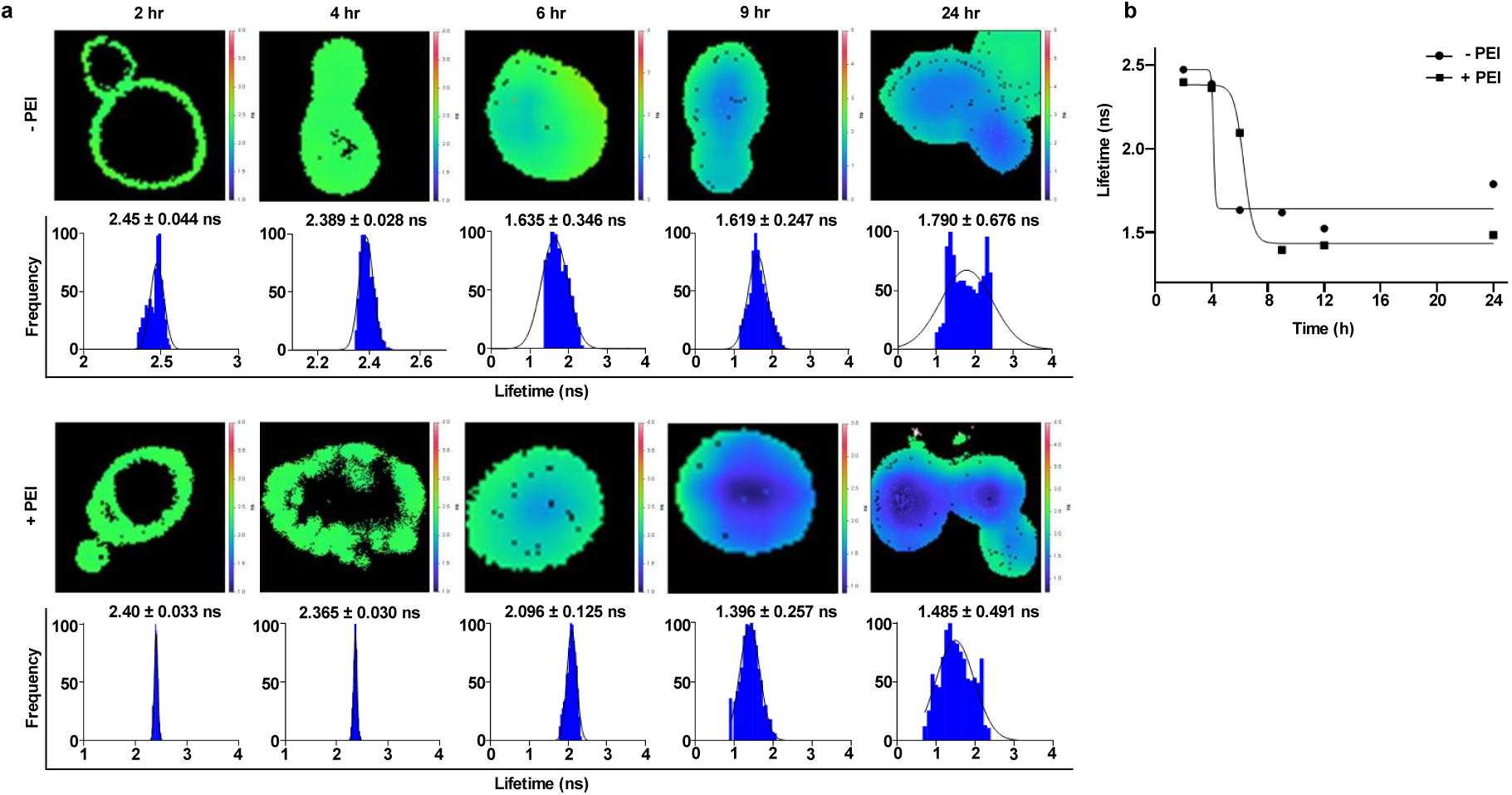
Lifetime analysis of GFP tagged α-syn protein in vivo: **a)** α-syn either in absence (top) or presence of PEI (bottom) were imaged and subjected to TD-FLIM analysis as described in methods section. Histogram below each image timepoint depicts the lifetime distribution after fitting the data by taking the first derivative of the rising of the decay **b)** data points from either non phase separated and phase separated pathways were plotted and fitted into Boltzmann sigmoidal equation

### LLPS increases toxicity of α-syn expressed in yeast

At first, we assessed the toxicity of PEI inside yeast cells, using a serial dilution spot assay (34). These experiments were performed after incubating the un-induced (without any α-syn expression) yeast cells in the absence and presence of different concentrations of PEI. We found that cells treated with at least up to 0.133 µM PEI showed growth similar to the untreated cells, which clearly suggest that the concentration of PEI used (0.133 µM) used in the present investigation did not have any significant effect on yeast cells (Fig. 3a). We then measured the colony forming units (CFU) using α-syn expressing strain to evaluate if and how the above mentioned (phase separation pathway, in the presence of PEI and non-phase separation pathway, in the absence of PEI) conditions affect cellular toxicity. These measurements were carried out at different time points (4h, 8h and 24 h), and we found that CFU in the presence of PEI was less at all time points, which indicate that the aggregation via phase separation pathway resulted in greater toxicity (Fig. 3b). We found the level of α-syn expression was found to be similar both in the absence and presence of PEI suggesting that PEI at 0.133 µM concentration does not hampers the protein synthesis machinery/pathways inside yeast cells, and PEI alone is not toxic (Fig. 3c, d).

We used FACS to quantify the toxicity inside yeast cells by employing CellRoxTM Deep Red (ThermoFisher Scientific, USA). It has been noted before that the increase in ROS can be proportional to the increased toxicity inside yeast (50). Cells in the absence and presence of PEI at different time intervals were treated with CellRoxTM Deep Red and subjected to FACS measurements. The percentage of cells showing ROS were plotted. For PEI treated cell population at each time interval there is a statistically significant increase in ROS production when compared to the untreated cell population further suggesting the increased toxic environment when α-syn undergoes aggregation via phase separation pathway (Fig. 3e).

Aggregates of proteins responsible for neurodegeneration are known to damage mitochondria of healthy cells (51). We imaged mitochondrial morphology to monitor the toxicity of the non-phase separation (in the absence of PEI) and phase separation (in the presence of PEI) pathways. At different time intervals, both untreated and PEI treated cells were incubated with MitoTracker Red and imaged under microscope. Healthy mitochondria in yeast appears as long intact tubular structure near the plasma membrane (52). We did not see any change in mitochondria after 4 hours. During the later time point, for example after 8 h, we observed increased fragmentation of mitochondria in PEI treated cells when compared with untreated cells. At 24 h there was complete degeneration of mitochondria in PEI treated cells but the untreated cells still had few visible mitochondria (Fig. 3f). To quantify this observation, we performed FACS using the same MitoTracker Red stained cells at 8 h and it was found that 87% of untreated cells were showing fluorescence corresponding to the presence of intact mitochondria whereas only 54% of cells treated with PEI showed intact mitochondria (Fig. 3g). We believe that, as the aggregates accumulate, they start binding directly to the membrane thereby altering the mitochondrial membrane potential ultimately leading to mitochondrial fragmentation (53, 54). Mitochondrial ROS was also measured at 8 h using MitoSOX RED dye which binds only to mitochondrial ROS species. Less than 1% of the untreated cells were seen to have accumulation of MitoSOX RED dye as compared with PEI treated cells where 42% of cells showed MitoSOX accumulation (Fig. 3h).

**Figure 3:**
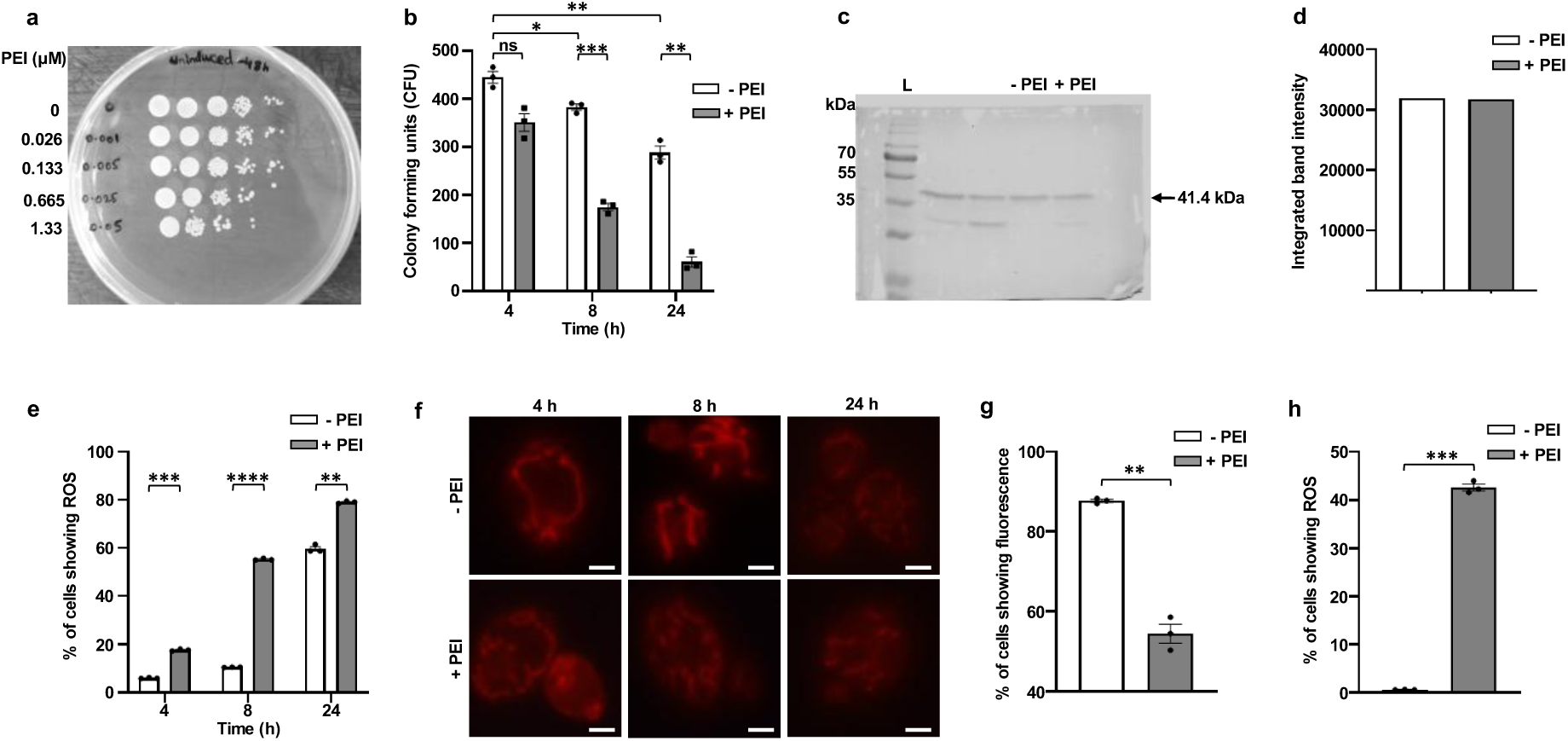
PEI treatment increases toxicity α-syn expressed in yeast: **a)** Serial dilution spot assay was performed for determining the toxicity of PEI on *S. cerevisiae* W303-1B cells. Cells were treated with different concentration of PEI for 48 h and spotted on SD agar plates and growth was assessed after 48 h at 30 °C (10-fold dilutions ranging from 10^1^ to 10^4^) **b)** PEI induced α-syn toxicity was determined by clonogenic assay. The number of colony-forming units (CFU) developed on SD agar plates at 30 °C for 72 h were recorded after treatment with PEI and counted using imageJ software **c)** Western blot of yeast cells expressing α-syn after induction for 8 h without and with PEI treatment was carried out by immunoblotting using anti-GFP antibody **d)** Intensity of bands on the membranes was quantified using imageJ software **e)** α-syn expressing yeast cells without and with PEI exposure for different time interval, the cells were stained with CellROX® Deep Red Reagent and analysed using flow cytometry **f)** MitoTracker Red fluorescence imaging of *S. cerevisiae* cells expressing WT α-syn-GFP in the absence and presence of PEI for different time intervals for visualization of mitochondrial fragmentation [Bar = 5 μm; magnification = 100× (oil immersion objective)] **g)** Flow cytometry quantitative analysis of mitochondrial morphology in yeast cells harbouring active mitochondria after treatment with PEI for 8 h **h)** After PEI exposure for 8 h, the cells were stained with MitoSOX RED and flow cytometric quantitative analysis was performed

### Aggregates characteristics differ between the phase separation and non-phase separation pathways

*S. cerevisiae* expressing α-syn incubated with and without PEI were grown in inducing media. After 8 h cells were lysed and subjected to dot blot analysis for the estimation of monomeric and fibrillar species of α-syn using specific antibodies (Fig. 4a). From the densitometric analysis, we found that, in the absence of PEI (under non-phase separation condition) a large fraction of α-syn existed as monomer with presence of small amount of fibrillar species or aggregates. In contrast, in the presence of PEI (under phase separation condition), the monomeric species gets drastically reduced and shows high concentration of fibrils (Fig. 4b). We then used proteinase K assay digestion to investigate the proteolytic fingerprints of the aggregates formed at the end of the phase separation (in the presence of PEI) and non-phase separation (in the absence of PEI) pathways. Aggregates formed under phase separation pathway were found to be more resistant to proteolytic digestion suggesting a more rigid and less solvent exposed structure as compared with aggregates formed in absence of phase separation pathway (Fig. 4c, d).

**Figure 4:**
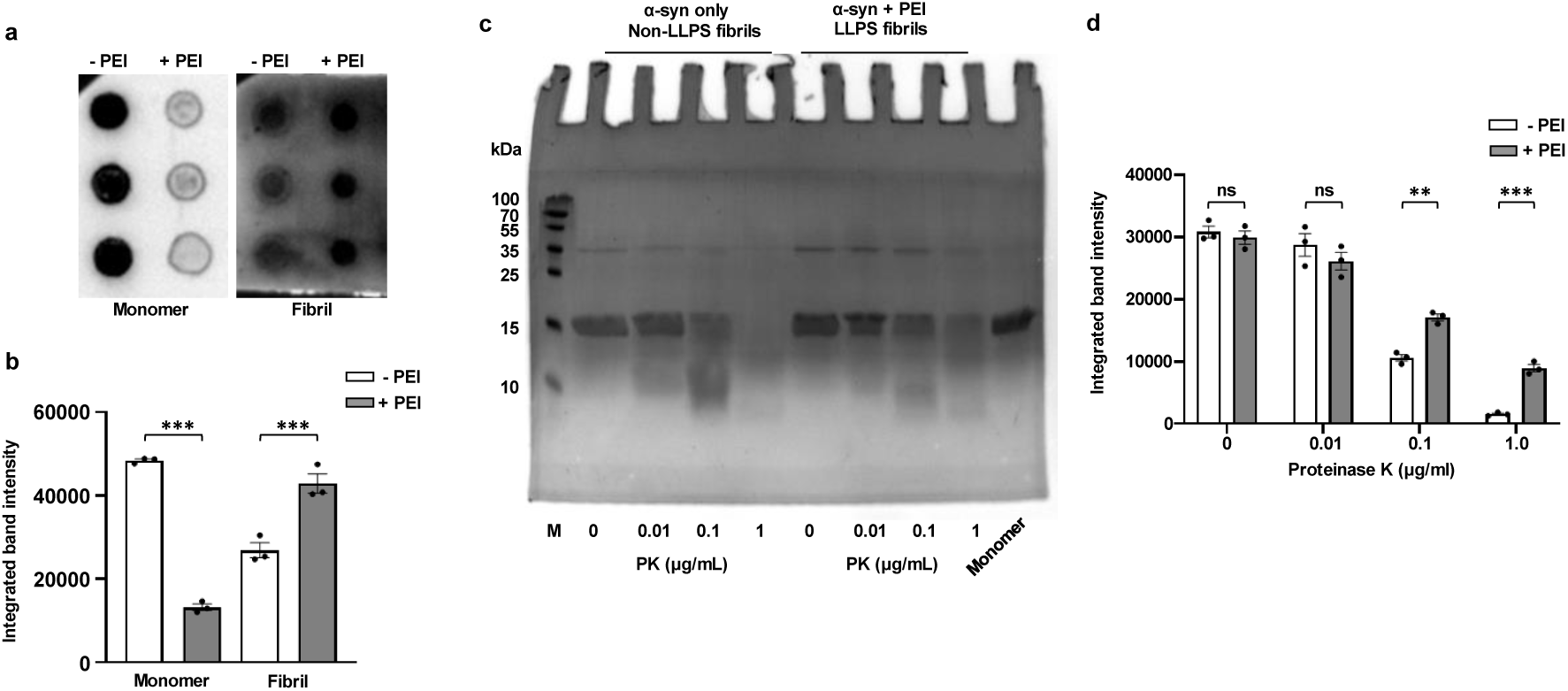
LLPS increases fibrillar load having distinct morphology: **a)** Dot blot analysis of yeast cell lysates expressing α-syn incubated without and with PEI for 8 h, membranes were probed using α-syn antibody specific for monomers and aggregates **b)** Intensity of dots on the membranes was quantified using imageJ software **c)** Proteolytic fingerprints of α-syn fibrils produced under LLPS and non-LLPS conditions were run on 15% SDS-PAGE gel **d)** Band intensities of the proteolytic fingerprints of α-syn under different conditions was quantified using imageJ

### In vitro experiments suggests that PEI binds to α-syn and facilitates phase separation

Subsequently we wanted to understand the mechanistic role of PEI on the phase separation of α-syn, and these experiments were carried out under in vitro conditions. First, docking study using AutoDock 4.2 predicted possible interactions between α-syn and PEI, with a dissociation constant (K_D_) of 1.97 nM (Supplementary Table 1). This suggests a strong binding affinity between positively charged PEI and negatively charged C-terminal domain of α-syn. We found further that glutamic and aspartic acid residues, namely, E110, D115, D119 and E123, were presumably involved in the interaction by forming hydrogen bonds along with additional polar contacts representing the favourable electrostatic interaction between the oppositely charged molecules (Supplementary fig. 3b). Further analysis of sequence properties using FuzPred (https://fuzpred.bio.unipd.it), which predicts context dependent binding behaviour of proteins (55, 56), suggested that the C-terminal region (102-140 aa) to have high propensity for multivalent interactions and to remain in the disordered form (Supplementary fig. 3c).

Fluorescence spectroscopy assay was used to measure the binding between α-syn and PEI, which suggested a strong binding with K_D_ of 15 nM when fitted into One site-total binding equation approaching saturation, which is within an order of magnitude of the computationally predicted value (1.97 nM) (Fig. 5a). When we incubated the protein in the presence of PEI, we found that α-syn underwent phase transition, which was reflected by measurable change in the turbidity of the solution at wavelength of 350 nm (absorbance value = 0.58) as compared with α-syn alone (Fig. 5b, supplementary fig. 4c). *In-vitro* droplet imaging also suggested that α-syn formed liquid droplets in the presence of PEI, but did not in its absence (Fig. 5c). Although the droplets of α-syn were dissolved on addition of 1,6-HD (10%), when they were allowed to mature, they did not dissolve. Instead, the matured droplets in the presence of 1,6-HD showed a prominent change in the circularity (Fig. 5c and d). This suggests that the droplets slowly undergo gelation, which was followed by the formation of aggregates as reported earlier (6). We then used 15% SDS page to calculate protein partitioning between the light (L) and dense phases (D) of the PEI induced droplets (Supplementary fig. 4d and e). Additionally, we probed how the far UV CD of α-syn change with time in the presence of different concentrations of PEI (Fig. 5e, supplementary fig. 5). With time, we found a decrease in the far UV CD characteristics, which became more prominent in the presence of PEI. When we plotted the apparent partition Gibbs free energy (Fig. 5f) and the ellipticity at 198nm at 24 hours, we found a qualitative similarity in their change, which seem to suggest that a conformational event in α-syn may have roles to play in the PEI induced phase transition of the protein (Fig. 5g).

**Figure 5:**
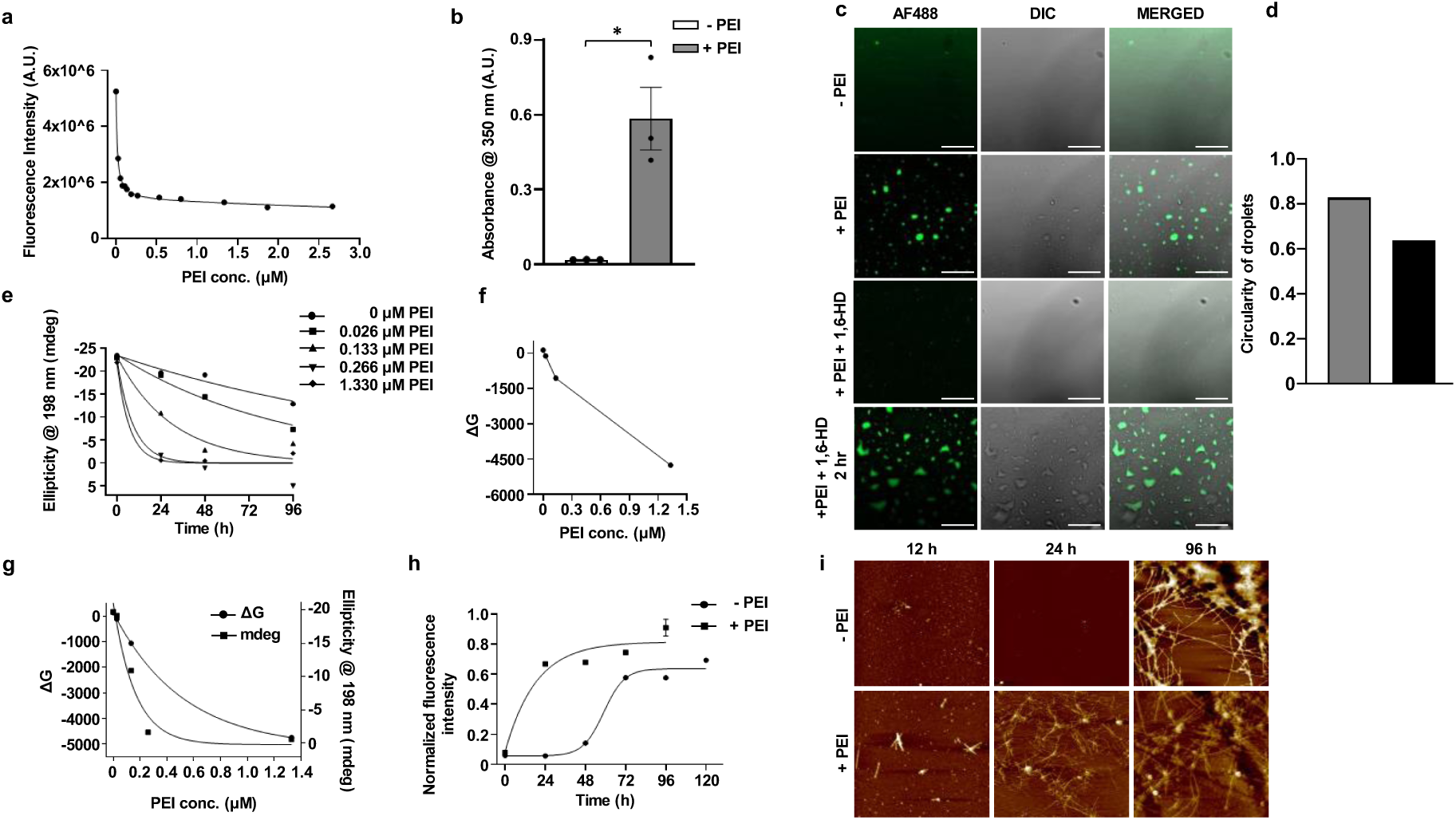
Binding of PEI to α-syn facilitates its phase separation and hence aggregation: **a)** Fluorometric binding analysis of α-syn protein upon titration with different concentration of PEI and fitted using non-linear one site binding equation gives a Kd value of 15.33 nM **b)** Turbidimetric analysis of α-syn protein upon interaction with PEI showing increased absorbance when measured at 350 nm owing to increased turbidity in the presence of PEI as compared with α-syn alone **c)** *In-vitro* droplet imaging of α-syn protein in the absence and presence of PEI showing droplets formation with PEI, which gets dissolved when treated with 1,6-HD (10%) [Bar = 20 μm; magnification = 63× (oil immersion objective)] **(d)** Matured droplets were not dissolved upon treatment with 1,6-HD; instead their circularity was changed which according to the previous literature, seemed to suggest the gelation of the droplets **e)** Circular dichroism spectra of α-syn protein at different time points treated with different concentration of PEI. The resulting curve was fitted into a single exponential growth equation **f)** Apparent partition Gibbs free energy from the dense and light phase formed by the treatment with different PEI concentrations incubated for 24 h **g)** Combined plot of the Apparent Partition Gibbs Free Energy and and the ellipticity value (in mdeg) at 24 h **h)** ThT-binding assay showing fibrillation of α-syn protein in the presence and absence of PEI at different time interval and was fitted into Boltzmann sigmoidal equation getting an aggregation specific curve showing fibril formation **i)** AFM images of α-syn protein in the presence and absence of PEI at different time interval showing fibril formation

### PEI induced LLPS of α-syn leads to its aggregation *in-vitro*

We then studied the aggregation kinetics of α-syn under phase separation and non-phase separation conditions. For both non-phase separation condition and phase separation condition, we incubated the protein at 37^0^C and studied different points using ThT fluorescence assay (29). We found that the protein under phase separation condition forms fibrils in a rate much faster than what was formed with α-syn alone (Fig. 5h). The fluorescence emission data were further complemented by atomic force microscopy, which was done using same samples used in ThT fluorescence assay. In the case of phase separation condition small proto-fibrillar structures starts appearing at about 12 h of incubation and a dense fibrillar network of α-syn aggregates can be seen after 24 h incubation which in case of non-phase separation condition appears only after 96 h of incubation (Fig. 5i).

## Discussion

While *in vitro* phase separation of intrinsically disordered proteins has been studied extensively, *in vivo* conditions are not yet clearly understood. *In vivo* systems are more complex, with many contributing factors, including the presence of multiple interaction partners, the effect of crowding environment and stress. The potential of using yeast as a model to study neurodegenerative diseases under *in vivo* condition has been established by others for α-syn in PD pathogenesis (18, 19). The present study aligns with the above approach while giving special emphasis to investigate the phase separation properties. We believe that this can be used as a simple yet effective system to investigate novel and/or repurposed small molecules, which can complement fast paced *in-vitro* studies.

Fig. 6 below outlines the data presented in this manuscript highlighting the presence of both the Phase Separation and Non-Phase Separation Pathways inside a yeast model. A balance between the long and short-range interactions is needed to form a dense phase in a droplet forming system (57). From the data presented here and from other previous literatures, it is apparent that PEI modulated phase separation of α-syn would be governed by the electrostatic interaction between the negatively charged C-terminal domain of α-syn and positively charged PEI. It is also interesting to note that the presence of PEI facilitates the phase separation and increases in the rate of aggregation, which is similar to the previous observation that PEI coated nanoparticles increased the aggregation of α-syn (46). Additionally, the aggregation kinetics under phase separation condition has been found to be hyperbolic, while that under non-phase phase separation condition is sigmoidal. This is expected for the seeded aggregation, as the droplets are known to contain early oligomeric molecules, acting as the seeds (6, 58, 59). Presumably, this is one of the reasons the increased rate of aggregation in the phase separation condition, although the exact mechanistic interpretation requires further study. In addition, we found that the basic characteristics of the fibrillar aggregates when formed under Phase Separation pathway vs under non-Phase Separation pathway. It may be noted that the heterogeneity at the fibril morphology has been observed in multiple studies, and these differences depend on various factors, including fibril forming conditions, the sources of the fibrils among others (60–70).

**Figure 6:**
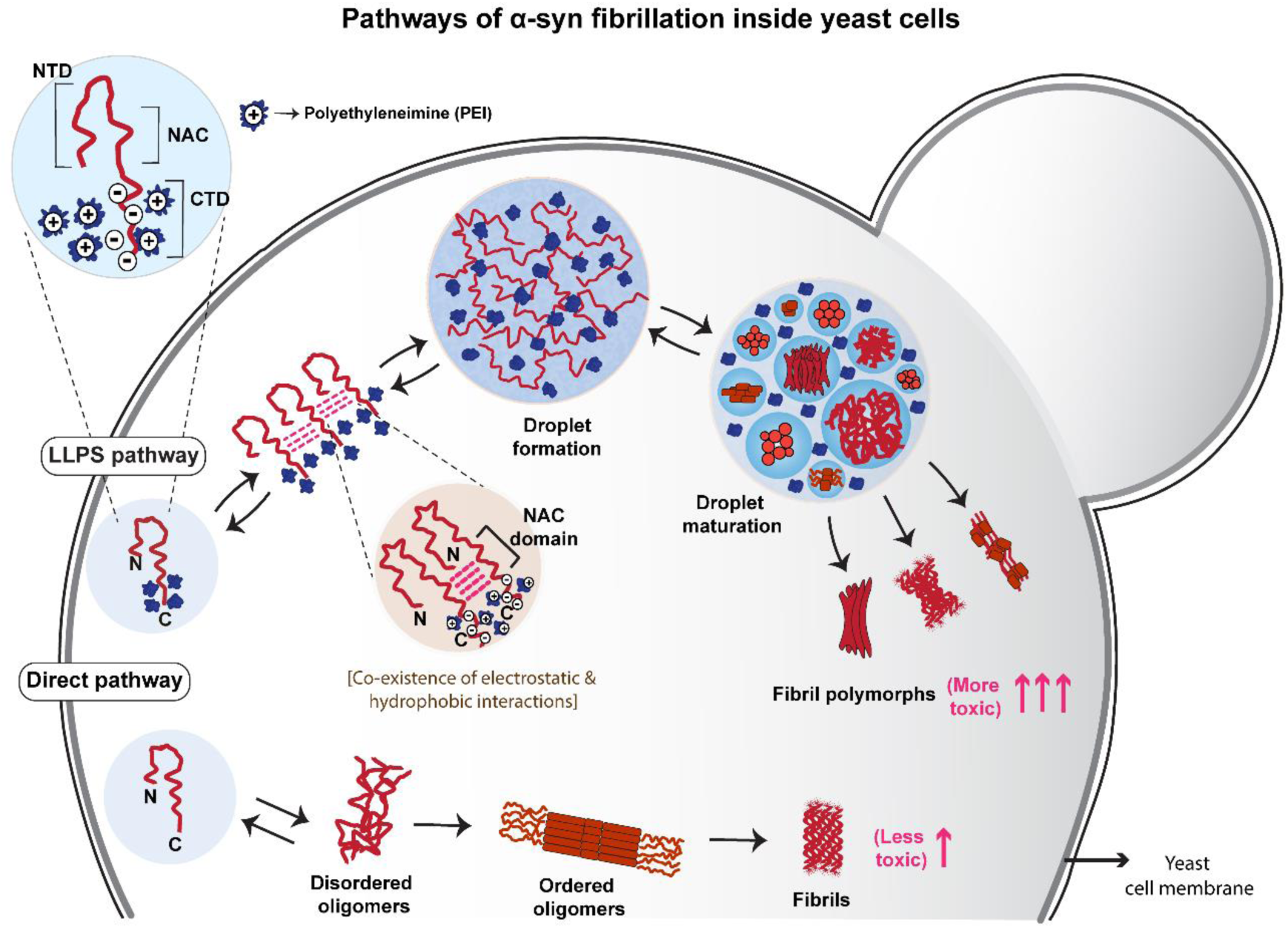
Proposed yeast model which suggests that α-syn aggregates formed via LLPS pathway are more toxic as compared with aggregates formed via non-LLPS pathway

The question of toxicity and its implications in the aggregation pathways in general is extensively debated. This has been further compounded by the question of what roles these proteins droplets might be playing in the heterogeneous landscape of protein aggregation. We have shown before using in vitro studies that protein droplets are characterized by significantly more toxicity than the late-stage fibrils (44). It is important to see that the present data inside yeast also support that observation. We are now studying the implications of this increased toxicity in neuro-degeneration and other roles other cellular machineries play inside yeast.

