## Supporting Information for "Multivalent interaction induces phase separation and formation of more toxic aggregates of α-syn in a yeast model of Parkinson’s disease"

### Supplementary Data

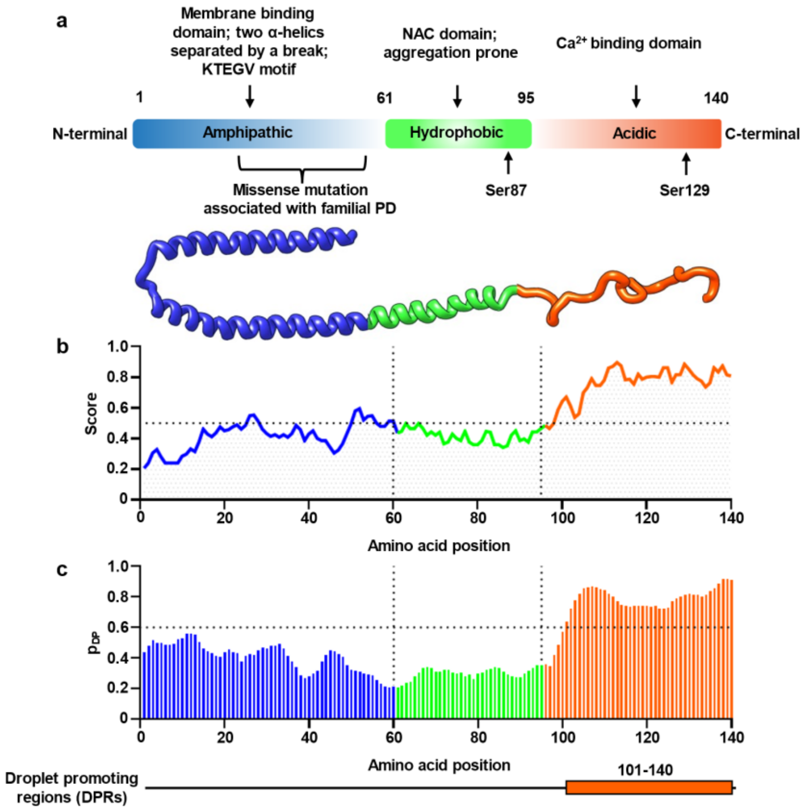


**Supplementary figure 1: *In silico* analysis of α-syn protein: a)** Structure of α-syn showing different regions and properties (blue amphipathic N-terminal, green hydrophobic NAC domain and orange acidic C-terminal) **b)** IUPred2A prediction of intrinsic disorder in α-syn **c)** FuzDrop algorithm showing the propensity of droplet promoting region based on sequence analysis

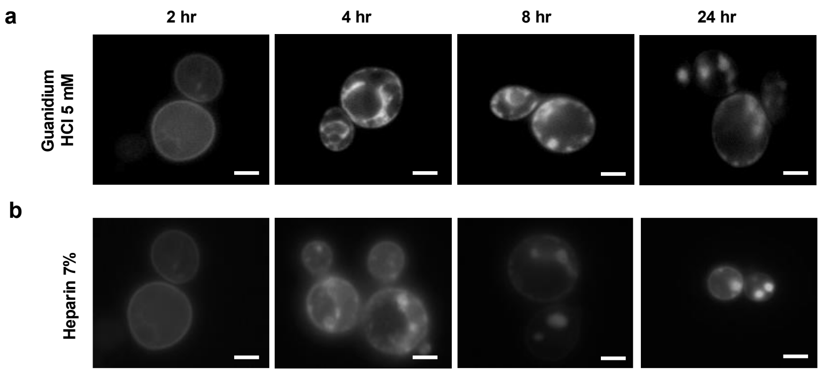


**Supplementary figure 2:** Fluorescence imaging of S. cerevisiae cells expressing WT α-syn-GFP in the presence of **a)** 5 mM Guanidium HCl, a known inhibitor of HSP104 **b)** anionic agent heparin (7%) [Bar = 5 µm; magnification = 100× (oil immersion objective)]

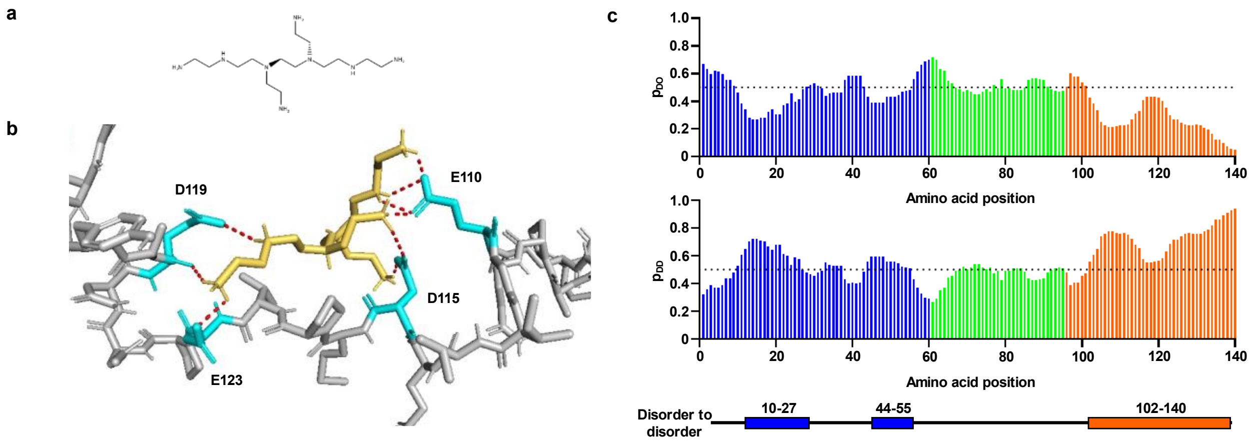


**Supplementary figure 3: *In-silico* binding analysis of α-syn with PEI: a)** Chemical structure of branched Polyethyleneimine (PEI) comprises of four $(C_2H_5N)_n$ monomers **b)** Polar interactions between α-syn and PEI representing the binding site comprises acidic residues E110, D115, D119, and E123 in C-terminus domain. Interacting residues were shown in cyan, ligand is in yellow, polar contacts were represented in red dotted lines and rest protein back bone in grey colour **c)** FuzPred algorithm predicting the context-dependent binding behaviour of proteins with regions of disorder-to-order probabilities ($p_{DO}$) and disorder-to-disorder probabilities ($p_{DD}$)

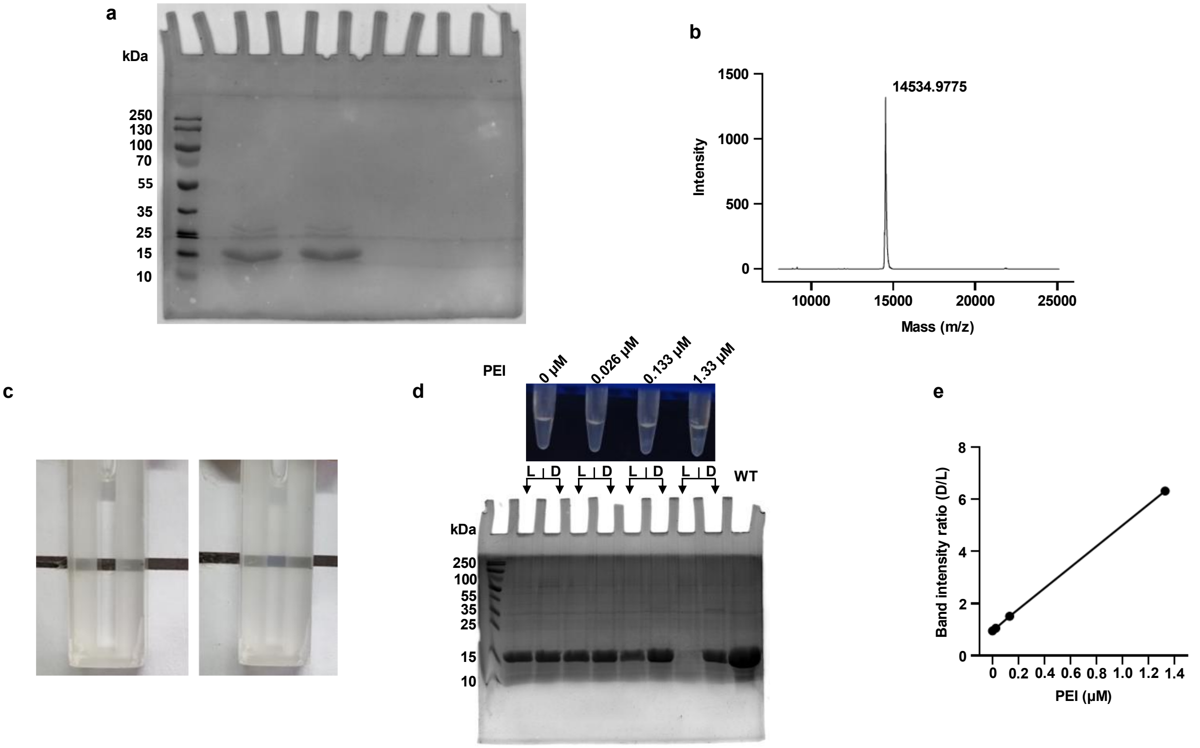


**Supplementary figure 4: a)** Purified α-syn protein was run on 15% SDS-PAGE showing single band at approx. 15 kDa corresponding to the theoretical molecular weight of wild type recombinant α-syn protein **b)** MALDI peak of purified monomeric α-syn protein at 14.5 kDa **c)** Cuvette showing visual difference in turbidity of α-syn protein upon interaction with PEI as compared with α-syn alone **d)** Fraction of protein from dense and light phase formed by treatment with different PEI concentration incubated for 24 h were run on SDS-PAGE (15%) **e)** Ratio of band intensity or gel partitioning (D/L) of dense phase (D) to light phase (L) was calculated and plotted

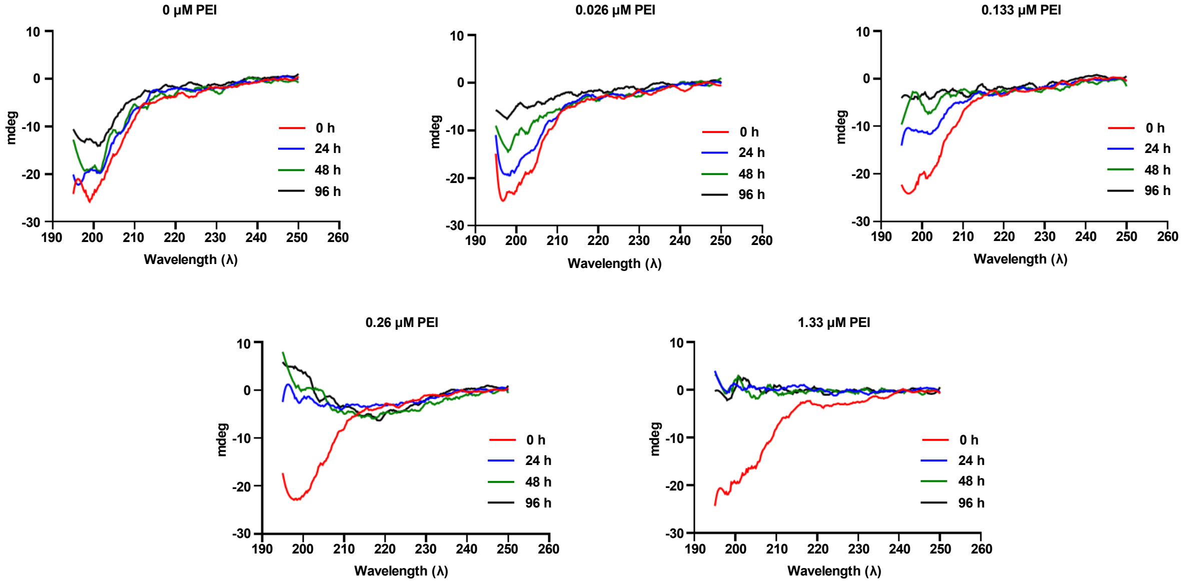

**Supplementary figure 5: Circular dichroism study of α-syn with PEI:** Change in the secondary structure of α-syn when treated with different concentrations of PEI was monitored by far UV-CD analysis at different time intervals

| | |
|---|---|
| Docking Energy (Kcal/mol) | -11.88 |
| Inhibition Constant ($K_i$) | 1.97 nM |
| Ligand efficiency | 0.54 |
| No. of H-bonds | 5 |

**Supplementary table 1: *In-silico* docking analysis of α-syn with PEI**
